# Audiovisual stimuli enhance narrowband gamma activity along the mouse thalamocortical visual circuit

**DOI:** 10.1101/2023.05.26.542476

**Authors:** Clément E. Lemercier, Patrik Krieger, Denise Manahan-Vaughan

**Affiliations:** Department of Systems Neuroscience; Department of Neurophysiology, Medical Faculty, Ruhr University Bochum, Bochum, Germany

## Abstract

To what extent thalamic activity can contribute to multisensory integration at cortical level is unclear. To explore this aspect, we used the mouse narrowband gamma oscillations (NBG), which arise from the lateral geniculate nucleus (LGN) and from upstream retinal inputs, as a tool to investigate potential thalamic audiovisual information transfer to the primary visual cortex (V1). We found that simultaneous bimodal audiovisual stimulation increased the power of V1 NBG. Pharmacological and optogenetic manipulations demonstrated that V1 NBG audiovisual responses occurred independently of primary auditory cortex activation. LGN recordings revealed that the majority of its neurons exhibited audiovisual properties. These properties comprised an increase of both the visual-evoked response and gamma-rhythmicity, indicating that the modulation of V1 NBG by audiovisual stimuli likely has a thalamic origin. Our results reveal a previously unreported subcortical source of audiovisual information transfer in V1 and suggest a new role for the LGN as a multisensory integration and relay center.

## Introduction

The last decade of research has provided fascinating insights as to how auditory information is integrated at the level of the primary visual cortex (V1) (Deneux et al., 2019; Garner et al., 2022; Ibrahim et al., 2016; Iurilli et al., 2012; Knöpfel et al., 2019; Majka et al., 2019; McClure and Polack, 2019; Meijer et al., 2018, 2020; Williams et al., 2023). In particular, by deciphering the functional role of crossmodal corticocortical auditory inputs existing from the primary auditory cortex (A1) to V1 (Deneux et al., 2019; Garner et al., 2022; Ibrahim et al., 2016; Iurilli et al., 2012), this research provided novel insights into context-dependent cortical neural circuits regulating V1 excitability, that serve to optimize crossmodal audiovisual information processing depending on changing ambient conditions (reviewed in Meijer et al., 2019). While the neural basis of audiovisual integration at the level of cortex become increasingly clear, basic questions as to whether crossmodal integration occurs at earlier stages along the sensory tracts are still largely unexplored.

In the recent years, it has emerged that some thalamic nuclei exhibit multisensory properties. For example, thalamic multisensory neurons have been identified in the auditory medial geniculate nucleus, that respond to visual, vestibular, and somatosensory stimuli (Komura et al., 2005). Furthermore, a subset of tactile-responsive neurons of the (somatosensory) ventral posteromedial (VPM) nucleus responds to visual stimuli (Allen et al., 2017), which in turn appear to strengthen thalamocortical coupling between the VPM and the primary somatosensory cortex (Bieler et al., 2018). Functional magnetic resonance imaging (fMRI) of human brain has indicated that the lateral geniculate nucleus (LGN), that comprises the first order visual thalamic nucleus, responds to auditory stimuli (Noesselt et al., 2010; Tyll et al., 2011), raising the intriguing possibility that the LGN is indeed multisensory, although its sensitivity to non-visual stimuli may be modality-specific (Bieler et al., 2018). The putative presence of audiovisual neurons in the LGN infers that they may play a role in integrating audiovisual information which then might be relayed at the level of V1. The challenge in addressing these intriguing possibilities derives from the difficulty in pinpointing thalamocortical communication that is not contaminated by crossmodal corticocortical A1-V1 inputs.

Recent studies have unveiled a novel narrowband gamma (NBG) oscillation that propagates and synchronizes throughout the mouse visual system (i.e. from the retina to high-order visual areas) (Koepsell et al., 2009; McAfee et al., 2018; Meneghetti et al., 2021; Saleem et al., 2017; Storchi et al., 2017; Shin et al., 2023), thereby coordinating the flow of visual information across visual areas (Shin et al., 2023). This type of oscillation is particularly driven by changes in light intensity and exhibits an inverse relationship with visual contrast (Meneghetti et al., 2021; Saleem et al., 2017; Storchi et al., 2017). This differs from the traditionally reported cortical broadband gamma (30-90 Hz) that is prompted by the receipt of visual contrast information derived from the synchronized activity of local excitatory-inhibitory circuits (Bastos et al., 2014; Buzsaki and Draguhn, 2004; Meneghetti et al., 2021; Saleem et al., 2017). At the level of V1, NBG is most strongly evident in the thalamo-recipient layer 4 (Meneghetti et al., 2022; Saleem et al., 2017; Shin et al., 2023) and is mediated by gamma-coherent inputs from the LGN (Saleem et al., 2017; Shin et al., 2023). The aforementioned characteristics make NBG an attractive model to investigate potential thalamic audiovisual information transfer to V1. We therefore recorded NBG from the mouse V1 during unimodal visual and bimodal audiovisual stimulation, under conditions of pharmacological and optogenetic manipulations of A1, and probed audiovisual properties of LGN neurons to clarify if the LGN can serve as a subcortical source of audiovisual information for V1.

## Material and methods

### Experimental model

Experiments were performed on wild type C57BL/6J mice (Charles River, Sulzfeld, DE), of both sexes, aged of 3-6 months. Experiments involving optogenetic manipulations were performed on mice that express channelrhodopsin selectively in glutamate acid decarboxylase-2 (GAD2)-positive neurons by crossing Ai32 (Ai32(RCL-ChR2(H134R)/EYFP; Strain #:012569, Jackson Laboratory, Bar Harbor, USA) and Gad2-IRES-cre (Strain #:010802, Jackson Laboratory) transgenic mice. Experimental procedures were conducted according to both European Union and German animal welfare regulations and were approved in advance by the local government ethics committee (Landesamt für Arbeitsschutz, Naturschutz, Umweltchutz und Verbraucherschutz, Nordrhein-Westfalen, Germany).

### Sensory stimulation

Visual stimulation was generated by means of a light emitting diode (diameter: 5 mm, 572 nm, 20°, 16,000 mcd) connected to a PVC-insulated polymethyl methacrylate (PMMA) optical fiber (outer diameter: 3.5 mm, inner diameter: 3 mm, length: 40 cm) controlled by a microcontroller (Arduino Uno Rev3; Arduino SRL, Monza, Italy). The tip of the optical fiber was placed ∼1 cm from the animal’s left eyeball and visual stimuli consisted of an iso-luminant full-field light flash of a 1 sec (∼100 lux, calibrated with TSL2561; Adafruit Industries, New York, NY, USA). Auditory stimulation was produced by a triggerable arbitrary waveform generator (Rigol DG811; Rigol Technologies, Inc., Portland, OR, USA), connected to a power amplifier (Nobsound AK170; Frequency response: 20 Hz – 20 kHz; Nobsound, Shenzhen Cavins Technology Co., Ltd., China) and a loudspeaker (FR 10 – 4 Ω; VISATON GmbH & Co. KG, Haan, Germany) placed at a distance of ∼30 cm to the left of the animal’s head. Auditory stimuli consisted of an open field white noise burst, delivered at various sound pressure levels (either 60-, 70– or 80-dB SPL, calibrated with Voltcraft SL-200; Conrad Electronic SE, Hirschau, Germany) using durations of either 1 sec, or 100 ms. Background noise level was ∼45-dB SPL. Stimulators were triggered by a programmable pulse generator (Master-9; A.M.P. Instruments, Ltd., Jerusalem, Israel).

### Imaging of intrinsic optical signals

The setup for intrinsic optical signal (IOS) imaging comprised a data acquisition system (Imager 3001, Optical Imaging, Inc., Rehovot, Israel), a CCD camera (Adimec 1000m; Adimec, Eindhoven, Netherlands) and a 90 mm macro lens (Tamron Co. Ltd., Saitama, Japan). Image resolution was 1,000 x 1,000 pixels for a pixel size of ∼10 µm. Illumination was generated by a halogen light source (Zeiss HAL 100; Carl Zeiss MicroImaging GmbH, Göttingen, Germany) under constant current supply, and illumination was time-controlled with an optical shutter (Uniblitz VS14; Vincent Associates, Rochester, NY, USA). Images of the superficial blood vessel pattern were obtained by focusing the camera on the pial surface, while illuminating the cortex at 546 nm. For functional imaging, the illumination wavelength was switched to 630 nm, focus was adjusted ∼400 μm below the brain surface, and prior to acquisition, light intensity was adjusted just below the camera’s saturation level. Data acquisition started after 2 sec of initial illumination and comprised 1 sec of baseline, 1 sec of stimulation and 4 sec of post-stimulation recordings. Sensory stimuli and data acquisition were synchronized by establishing communication between the Imager 3001, the optical shutter, and the programmable pulse stimulator (Master-9). Raw images were acquired at a rate of 50 frames per second (fps) and then temporally binned to a rate of 5 fps. Change in reflectance relative to baseline (ΔR) was computed with the following equation: ΔR=(R-R_0_), whereby ‘R’ is amount of light reflected during any given frame and ‘R_0_’ is the average amount of light reflected prior start of the sensory stimulation. To improve the signal-to-noise ratio, a minimum of 3 trials, with 20 sec intertrial intervals, were averaged. Cortical responses to sensory stimuli were visualized by using the WinMix 1.9 software (Optical imaging Inc.) after spatial binning (10×10) and clipping (generally 2-3 times the image standard deviation).

### Surgical and electrophysiological procedures

Mice were anesthetized by an intraperitoneal injection of a physiological saline mixture of urethane (1.2–1.4 g/kg, Sigma-Aldrich Chemie GmbH, Steinheim, Germany) and acepromazine (0.5 mg/kg, CP-Pharma Handelsgesellschaft mbH, Burgdorf, Germany). Body temperature was maintained at 37°C with a closed loop heating pad (FHC Inc., Bowdoin, ME, USA), oxygen (≥ 99,5 %) was supplied continuously, and the animal’s breathing rate was monitored on an oscilloscope by means of a piezoelectric disc (27 mm diameter) placed beneath the animal’s torso (Zehendner et al., 2013). The scalp was locally anesthetized with bupivacaine (0.25 %, mibe GmbH Arzneimittel, Brehna, Germany), incised, and a custom head-fixation implant comprising an attachment point and a circular hole used as recording chamber was fixed onto the skull with dental acrylic (Paladur, Kulzer GmbH, Hanau, Germany). The skull was gently thinned to transparency with a dental drill (OS-40; Osada Electric Co., Ltd. Tokyo, Japan) and IOS imaging was then performed to functionally guide the craniotomies (∼1 mm²) over V1 and A1. In another set of experiments, a craniotomy of ∼1 mm^2^ was performed over the LGN, at 2.3 mm lateral and 2.5 mm posterior to bregma, as described by others (Román Rosón et al., 2019).

Intracortical local field potentials (LFPs) were obtained from ∼1.5 MΩ borosilicate microelectrodes inserted in the layer 4 of V1 and A1 with the help of two separate stereotaxic micromanipulators (SM-25C; Narishige Scientific Instruments, Tokyo, Japan). Signals were amplified (NL100AK, NL104; Digitimer Ltd., Welwyn Garden City, UK), high-pass filtered at 0.1 Hz, digitalized at 10 kHz (Axon Digidata 1550 B; Molecular Devices, LLC. San Jose, CA, USA) and visualized with the pClamp 11 software (Molecular Devices).

Single-unit juxtacellular recordings of LGN neurons were obtained using 2-3 MΩ borosilicate microelectrodes with the help of a motorized micromanipulator (SM-1; Luigs & Neumann GmbH, Ratingen, Germany). Signals were amplified (AxoClamp 2B; Molecular Devices), high pass filtered at 300 Hz, digitalized at 20 kHz (Axon Digidata 1320 B; Molecular Devices), and visualized with the pClamp 8 software (Molecular Devices). In both cortical and thalamic recordings, glass microelectrodes were filled with an extracellular solution containing in mM:135 NaCl, 5.4 KCl, 1.8 CaCl_2_, 1 MgCl_2_, and 5 HEPES (pH ∼7.2).

### Imaging of the pupil

Images of the pupil were acquired under infrared illumination at 40 fps with the Raspberry Pi HQ camera module (no IR-cut filter; Raspberry Pi Foundation, Cambridge, UK) mounted on a 16 mm telephoto lens (CGL Electronics Co. Ltd., Hong Kong, China). Frames were cropped around the pupil (width: 160, height: 90 pixels) and extracted with the FFmpeg program (Tomar, 2006). Pupil segmentation and tracking of its diameter were performed with the ImageJ software (NIH, Bethesda, MD, USA).

### Ligand infusion

Activity of A1 was silenced with an infusion of ∼1 µL of the GABA_A_-receptor agonist, muscimol (Tocris Bioscience, Wiesbaden-Nordenstadt, Germany), at a dose of 1 mM dissolved in extracellular solution. The mixture was loaded in the recording glass microelectrode and was ejected by applying gentle positive pressure (Inverted suction-pulser; Sigmann Elektronik GmbH, Hüffenhardt, German). To allow the mixture to flow out of the electrode, electrode tip was mechanically enlarged. As a result, electrode resistance decreased from ∼1.5 MΩ to ∼0.2 MΩ and auditory evoked potentials exhibited smaller amplitudes than when recorded with regular electrodes.

### Optogenetic stimulation

Photostimulation of GAD2-expressing neurons via channelrhodopsin was achieved with 470 nm light pulses delivered by a fiber-coupled LED light source (M470F1; Thorlabs, Newton, NJ, USA). The pulse length was 1.1 sec, starting 100 ms prior to sensory stimulation, with a light intensity ranging from 0.6 to 2.4 mW. The output power of the LED driver (DC2100; Thorlabs) was controlled by the voltage output of the programmable pulse stimulator (Master-9). The power output at the optic fiber cannula (CFM52L10; Thorlabs) tip was measured with a photodiode power sensor (PM100D; Thorlabs).

### Recording Protocol

Stimulus presentation was not randomized, and recording blocks comprised 20 trials, repeated with an interstimulus interval of 10 sec. In all experiments, audiovisual blocks were interleaved by control blocks comprising visual stimulation alone.

### Spectral analysis of local field potentials

LFP time-series were down-sampled to 1 kHz and bandpass filtered between 0.1 to 300 Hz, with the R package ‘signal’ (Signal developers, 2013). The wavelet power spectrum of the filtered LFP time-series was obtained by applying a Morlet wavelet transform with the R package ‘WaveletComp v.1.1’ (Roesch and Schmidbauer, 2018). The detection period range was set from 8 to 1024 ms, in which each octave was divided into 60 suboctaves. In accordance with the approach described in Berens et al., (2008), V1 LFP time-series were divided into three different time periods: **1**. A baseline period of 800 ms prior to stimulus onset containing the spontaneous LFP fluctuations; **2**. A transient a period of 200 ms after stimulation onset, containing the initial event-related change in the LFP (i.e. visual-evoked potential); and **3**. a stimulation period ranging from 200 ms to 1 sec after stimulation onset, containing later event-related changes in the LFP (i.e. visual-evoked oscillatory activity). The power increase (Δ Power) in the LFP during the stimulation period (stim) relative to the baseline period (base), was defined by subtracting the power spectral density values of these two phases (Δ Power = Power_stim_ – Power_base_). From these differential spectra, power and frequency of the oscillations were quantified by measuring either their peak, or band power (40-to-80 Hz), and peak frequency.

### Analysis of single-unit activity

Spike detection was conducted using the threshold search function of the software pClamp 11.2 (Molecular Devices). Spike trains were divided into two phases, a baseline phase of 1 sec immediately prior stimulation onset, and a stimulation phase of 1 sec in which the sensory stimuli were delivered. Cut-off value for significant variation in spike rate was set at ± 1 Hz. Spike train power spectra were computed from binary spike train data with the R package ‘psd’ (Barbour and Parker, 2014).

### Statistics

Statistical analysis was performed using the R software (R Core Team 2020). Unless stated, data in the text are reported as M (IQR); where ‘M’ is the median and ‘IQR’ is the interquartile range. In the results section, ‘N’ corresponds to the number of animals and ‘n’ corresponds to the number of neurons. Differences between matched and unmatched measurements were respectively evaluated with the Wilcoxon signed-rank test (Figure 2 and Figure 4) and Wilcoxon-Mann-Whitney U test (Figure 3). Hypotheses were two-sided and tested at a level of significance of *p*< 0.05. To report statistics in the text, visual and audiovisual conditions are respectively abbreviated as V and AV. Strength and direction of monotonic association between two variables were assessed with the Spearman’s rank correlation coefficient (Figure 4). Figures were produced with the R software ‘base’ package and ‘ggplot2’ package (Wickham 2016).

## Results

### V1 exhibits narrowband gamma and cross-modal auditory responses

First, we explored if we could detect V1 narrowband gamma oscillations (NBG) with intracortical local field potential (LFP) recording from the layer 4 (Figure 1A-1C). We directed our attention towards this layer because of the prominence of visual-induced NBG in this layer (Meneghetti et al., 2022; Saleem et al., 2017; Shin et al., 2023; Welle and Contreras, 2016). To avoid intentional movements in response to sound (Yeomans and Frankland, 1995), which have been reported to activate V1 (Bimbard et al., 2023; Williams et al., 2023), the recordings were conducted under anaesthesia. In response to an iso-luminant LED flash, V1 LFP power spectra exhibited sharp peaks in the gamma-frequency range (Figure 1C). These oscillations had a peak frequency of 69.1 Hz (2.5) and a relatively narrow half-power bandwidth (58.6 (2.5) – 81.8 Hz (0.7)) (N = 9) which is consistent with the characteristics of V1 NBG reported in other studies (McAfee et al., 2018; Meneghetti et al., 2021; Saleem et al., 2017; Shin et al., 2023; Storchi et al., 2017). In contrast to V1, the LFP power spectrum of A1 in response to white noise (60-to-80-dB SPL) did not exhibit rhythmical fluctuations, but rather comprised ‘ON’ and ‘OFF’ auditory responses (Figure 1C). The early sensory response (i.e., the evoked potential) in V1 and A1 were of comparable amplitudes (V1 = –0.28 mV (0.19), A1 = –0.36 mV (0.45); *p* = 0.5781, N = 7; Figure S1A and S1B), but the auditory response in A1 occurred twice as fast as the visual response in V1 (A1 = 23 ms (3.5), V1 = 45 ms (8.5); *p* = 0.0156, N = 7; Figure S1A and S1B).

**Figure. 1.**
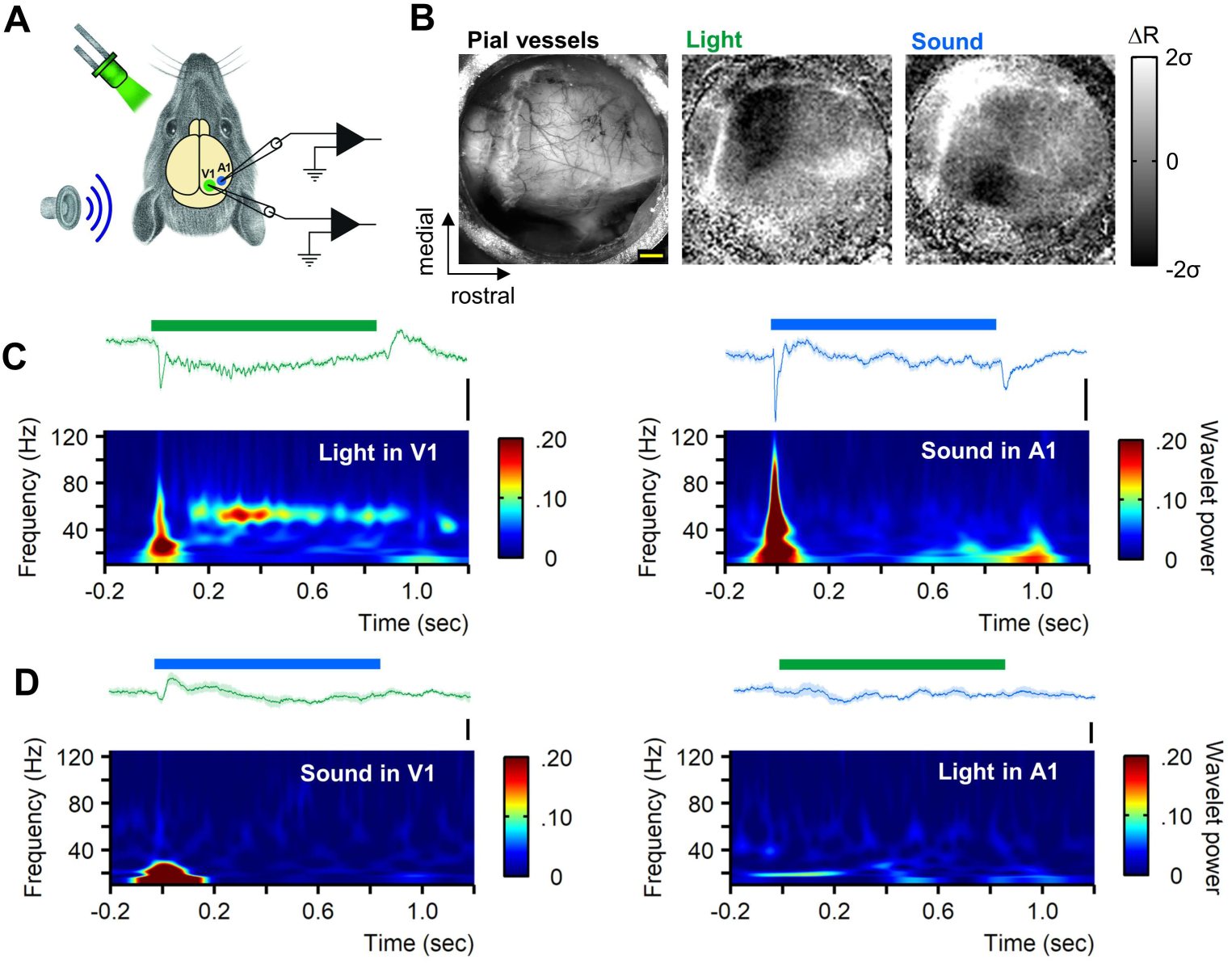
V1 exhibits narrowband gamma activity and cross-modal auditory response. (**A**) Illustration of the recording setup and stimulators used to deliver visual and auditory stimuli. **(B)** Example of functional delineation of V1 and A1 with IOS imaging. From left to right: background image of the skull surface and pial vessels (illuminated at 546 nm); average change (over 4 trials) in cortical light reflectance (illuminated at 630 nm) in response to visual (1 sec iso-luminant LED flash at ∼100 lux) and auditory stimuli (1 sec white noise at ∼80-dB SPL). Response frames for visual and auditory stimuli were selected respectively 2 and 1 sec after stimulation onset (data are from the same animal). Scale bar is 1 mm; scale and orientation scale apply to all images. **(C)** Examples of LFP traces and corresponding wavelet power spectra showing typical visual and auditory responses in V1 (left) and in A1 (right) (average of 20 trials, data from the same animal; scale bars: 0.4 mV. **(D)** Grand mean LFP traces (average of 130 trials, N = 7) and corresponding wavelet power spectra showing the presence of cross-modal auditory responses in V1 (left) and the absence of a cross-modal visual response in A1 (right). Scale bars: 0.1 mV.

We then investigated V1 and A1 cross-modal sensory responses. Consistent with previous reports (Iurilli et al., 2012), V1 exhibited cross-modal auditory responses pertained in the LFP in the form of a transient upward deflection in response to sound (Figure 1D). This response was described as relying upon direct corticocortical connections from A1 to V1 and thought to reflect local activation of GABAergic inhibition of V1 supragranular layer (Iurilli et al., 2012). In our experiments, this response had a latency to peak of 77 ms (110) and reached 0.14 mV (0.11) (130 trials, N = 7, Figure S1C and S1D). In contrast to V1, A1 LFPs did not exhibit cross-modal visual responses (Figure 1D). This indicates that auditory information is relayed to V1 but that reciprocal transmission of visual information to A1 could not be evidenced in our conditions, although it was reported that visual stimuli can modulate A1 activity (Kayser et al., 2008; Morrill and Hasenstaub, 2018; Wallace et al., 2004).

### Audiovisual stimuli increase the power of V1 NBG

Since V1 exhibited a cross-modal auditory response (Figure 1D), we then scrutinized the effects of bimodal audiovisual stimuli on V1 activity in more detail. We compared V1-evoked activity during unimodal visual and bimodal audiovisual stimulation and found that the power of V1 NBG was remarkably increased by audiovisual stimuli (Figure 2A). The effect of sound was pressure-dependent: the abovementioned power increase was evident when auditory stimuli were delivered at 70-dB (*p* = 0.0312, N = 7), and 80-dB (*p* = 0.0078, N = 9), but not when the sound pressure level was reduced to 60-dB SPL (*p* = 0.8438, N = 8; Figure 2B and 2C). Strikingly, sound delivered at 60dB continued to generate an auditory response in A1 (Figure 2C), albeit of lower amplitude. This suggests that the integration of auditory information through an increase in V1 NBG power is contingent on sound saliency i.e., only triggered by loud auditory stimuli. The oscillatory frequency of V1 NBG remained unchanged in response to audiovisual stimuli (*p* = 0.4412, N = 9; dataset exposed to 80-dB SPL). Examining V1 NBG power across audiovisual trials revealed a sudden increase in power in response to the first audiovisual stimulus, which then remained stable for the rest of the trials (Figure S2A and S2B).

**Figure 2.**
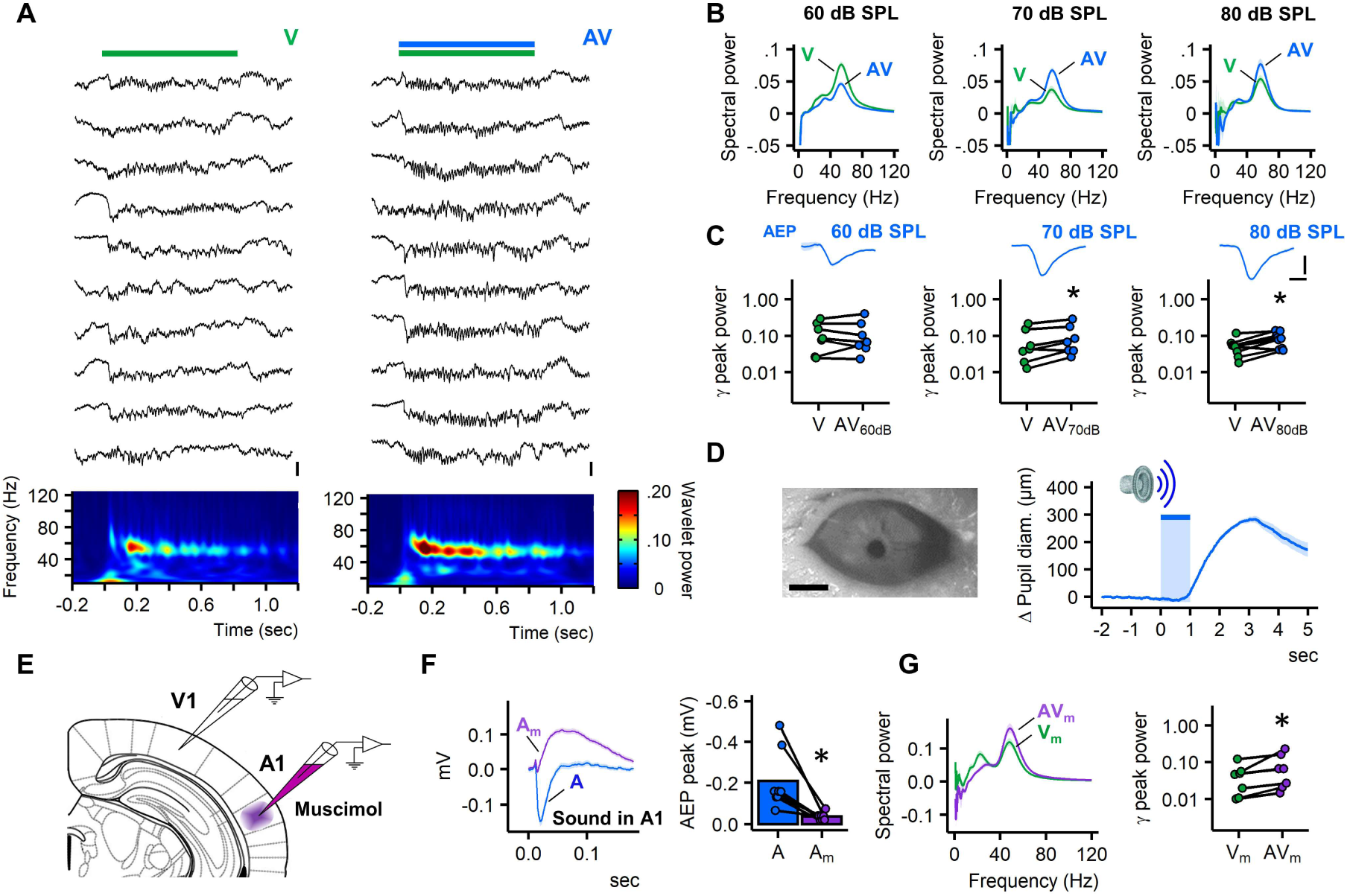
Audiovisual stimuli increase the power of V1 NBG independently of change in pupil size or A1 activation. (**A**) Example of 10 consecutive LFP traces and corresponding wavelet power spectra showing V1 responses to visual (V) and audiovisual stimuli (AV, 80-dB SPL); data from the same animal, scale bars: 0.4 mV. **(B)** Example of V1 response power spectra during V and AV stimuli (average of 20 trials, spectra at various sound pressure levels (SPL) were selected from different animals). (**C**) Quantification of V1 NBG peak power for V and AV stimuli provided at various SPL (60-, N = 8; 70-, N = 7; 80-dB SPL, N = 10). Above are shown example LFP traces of AEPs at the three different tested SPL (average of 10 trials, data from the same animal; scale bars: (y): 0.2 mV; (x): 10 ms). **(D)** Infrared photograph of the mouse pupil; effect of sound (80-dB SPL) on the mouse pupil diameter (average of 20 trials, data selected from one experiment out of a total of two performed; scale bar: 1 mm). **(E)** Illustration of the setup used for the experiments with local infusion of muscimol in A1 (Franklin and Paxinos, 2008; Plate 54). **(F)** Average auditory response before (blue) and after (purple) infusion of muscimol (∼1 µL at 1mM; average of 80 trials, N = 8); quantification of the auditory response peak amplitude before and after infusion of muscimol (N = 8). **(G)** Example V1 response power spectra during V and AV stimuli in condition of silenced A1 activity (average of 20 trials, data from the same animal); quantification of V1 NBG peak power during V and AV stimuli in condition of silenced A1 activity (80-dB SPL, N = 8). In (F-G), the letter ‘m’ denotes the muscimol condition. * Indicate *p* < 0.05; Wilcoxon signed-rank test.

After cessation of audiovisual stimuli, the power of V1 NBG returned to levels comparable to those of the baseline, although a few subsequent trials continued to exhibit significantly higher oscillatory power (Figure S2A and S2B).

Sound induces pupil dilation (Deneux et al., 2018; Montes-Lourido et al., 2021), and this may have triggered the change in NBG power detected in V1 in response to audiovisual stimuli. Measurements of pupil diameter in response to sound indicated that pupil dilation commenced ∼1 sec after the V1 sensory response (Figure 2D). Hence, the observed increase of V1 NBG power by audiovisual stimuli cannot be attributed to a greater amount of light passing through the pupil.

Next, we investigated whether the audiovisual-induced increase in V1 NBG power required the activation of A1. We significantly suppressed the auditory response in A1 by pharmacologically activating GABA_A_ receptors by means of local application of muscimol to A1 (*p*= 0.0078, N = 8; Figure 2E and 2F). The increase in V1 NBG power in response to audiovisual stimuli was unaffected after local inhibition of A1 (*p* = 0.0156, N = 7; Figure 2G). This finding was further supported by a comparable increase in V1 NBG power to that observed in the control experiments (AV_muscimol_ = +41,9 % (33), N = 7; AV_control_ = +71.6 % (78.5), N = 8; *p*= 0.6065). We verified that A1 did not contribute to the V1 NBG audiovisual response by optogenetically inhibiting A1 auditory response with local photo-activation of channelrhodopsin in Gad2– expressing neurons (Figure S3). Taken together, these results indicate that activation of A1 is not required to trigger the V1 NBG audiovisual response, which in turn implies that this effect is not mediated by corticocortical communication from A1 to V1.

### Audiovisual congruency influences V1 NBG response

We then aimed to understand to which extent V1 NBG audiovisual response depends on audiovisual temporal binding. To this end, various stimulation paradigms with variable sound durations and timings were implemented (Figure 3). For control purposes, we analyzed the differential wavelet power spectra between two blocks consisting solely of visual stimuli (Figure 3A). These spectra exhibited consistent oscillatory power across trials (Figure 3B). Fully temporally synchronized audiovisual stimuli increased the power of V1 NBG over the entire duration of the stimulation period (*p* = 0.0104, N = 8; Figure 3C). A comparable increase in the power of V1 NBG occurred when audiovisual stimuli were delivered for only 100 ms at the start of the visual stimulation (*p* = 0.0359, N = 9; Figure 3D). The delivery of an auditory stimulus for 100 ms, with a latency of 100 ms before initiating the visual stimulation, also resulted in an enhancement of V1 NBG power (*p* = 0.0379, N = 8; Figure 3E), but effects were rather transient, primarily occurring during the initial phase of the visual oscillatory period.

**Figure 3.**
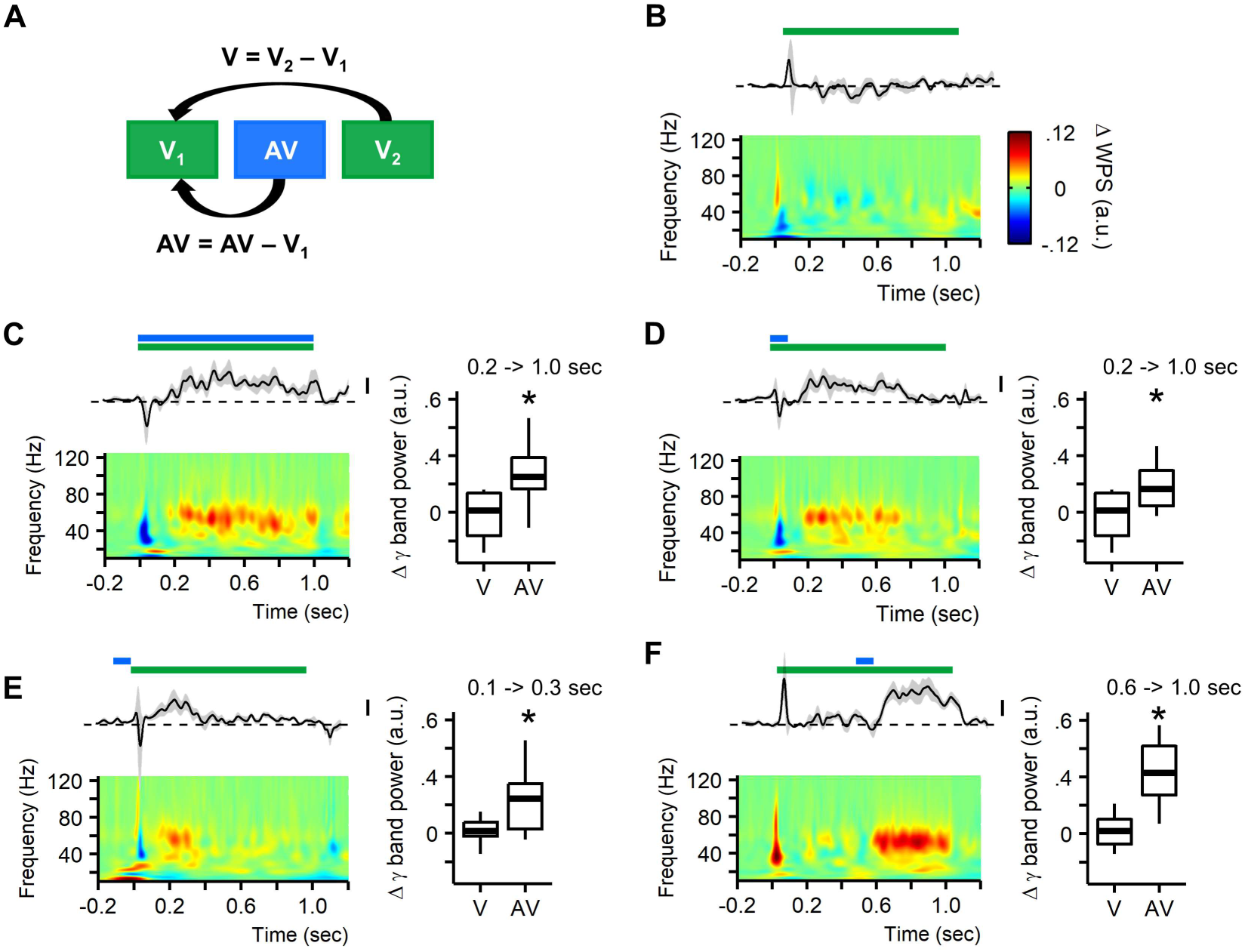
Effect of audiovisual congruency on V1 NBG response. (**A**) Illustration of the method used to compute differential wavelet power spectra (ΔWPS). **(B)** Variation in the power of V1 NBG across two blocks of trials with visual stimuli alone (V2 – V1). **(C-F)** Variation in the power of V1 NBG in response to (C) fully matched (N = 8), (D) partially matched (N = 9), (E) and unmatched (N = 8) audiovisual stimuli, as well as (F) brief auditory stimulation delivered in the middle of the visual stimulation (scale bars: 2/100 a.u.). Above each spectrum the average variation over time in NBG-band power (40-to-80 Hz band power) is shown. In each panel, boxplots show the change in NBG-band power during audiovisual condition (AV) relative to control (V, N = 8) as described in panel in (A). The reported time ranges indicate the interval of time in which the change in NBG band power is significantly different to control (Visual condition alone, panel B). In boxplots (C-E), * indicate *p* < 0.05; Wilcoxon-Mann-Whitney U test.

Remarkably, delivering an auditory stimulus of 100 ms at the midpoint of a visual stimulation resulted in a dramatic increase of the power of V1 NBG that persisted for the remainder of the subsequent visual stimulation period (*p* = 0.0047, N = 6; Figure 3F). Another interesting observation from this stimulation paradigm was that similarly to the effect of sound alone on V1 LFP (Figure 1D), the subsequent increase in V1 NBG power in response to sound was frequently preceded by an upward deflection in V1 LFP (Figure S4A and S4B), suggesting that prior to exerting its enhancing effect, sound induces a transient inhibition of V1 NBG. These findings indicate that a brief elevation in sound pressure level can significantly increase the power of V1 NBG, but also that V1 exhibits a relatively wide temporal window (of at least a hundred milliseconds) within which it can integrate auditory stimuli. Moreover, V1 NBG audiovisual response appears to exhibit a bimodal progression comprising an initial transient desynchronization followed by an increase in synchrony.

### LGN neurons exhibit audiovisual properties consistent with the increase of V1 NBG power

Since the audiovisual modulation of V1 NBG appears to be independent of A1 activation (Figure 2F-2G and Figure S3) and given that recent research suggests the LGN to be the primary source of V1 NBG (Meneghetti et al., 2021; McAfee et al., 2018; Saleem et al., 2017; Shin et al., 2013), we next explored whether this effect could have a thalamic origin. To test this possibility, we conducted juxtacellular recordings in the LGN during unimodal visual and bimodal audiovisual stimulation (Figure 4). LGN neurons were identified by their rapid firing in response to either switching-on (LGN-ON neurons) or switching-off (LGN-OFF neurons) light flashes delivered at the contralateral visual field of the animal (Piscopo et al., 2013) (Figure S5A). At the level of the recorded population (recording depth: 2575 μm (372), n = 40; Figure S5B), audiovisual stimuli significantly increased the spike rate of LGN-ON neurons (V = 11.77 Hz (11.46), AV = 13.02 Hz (13.92); *p* = 0.0016, n = 40, N= 10; Figure 4A). Audiovisual modulation of LGN-OFF neurons was not investigated.

**Figure 4.**
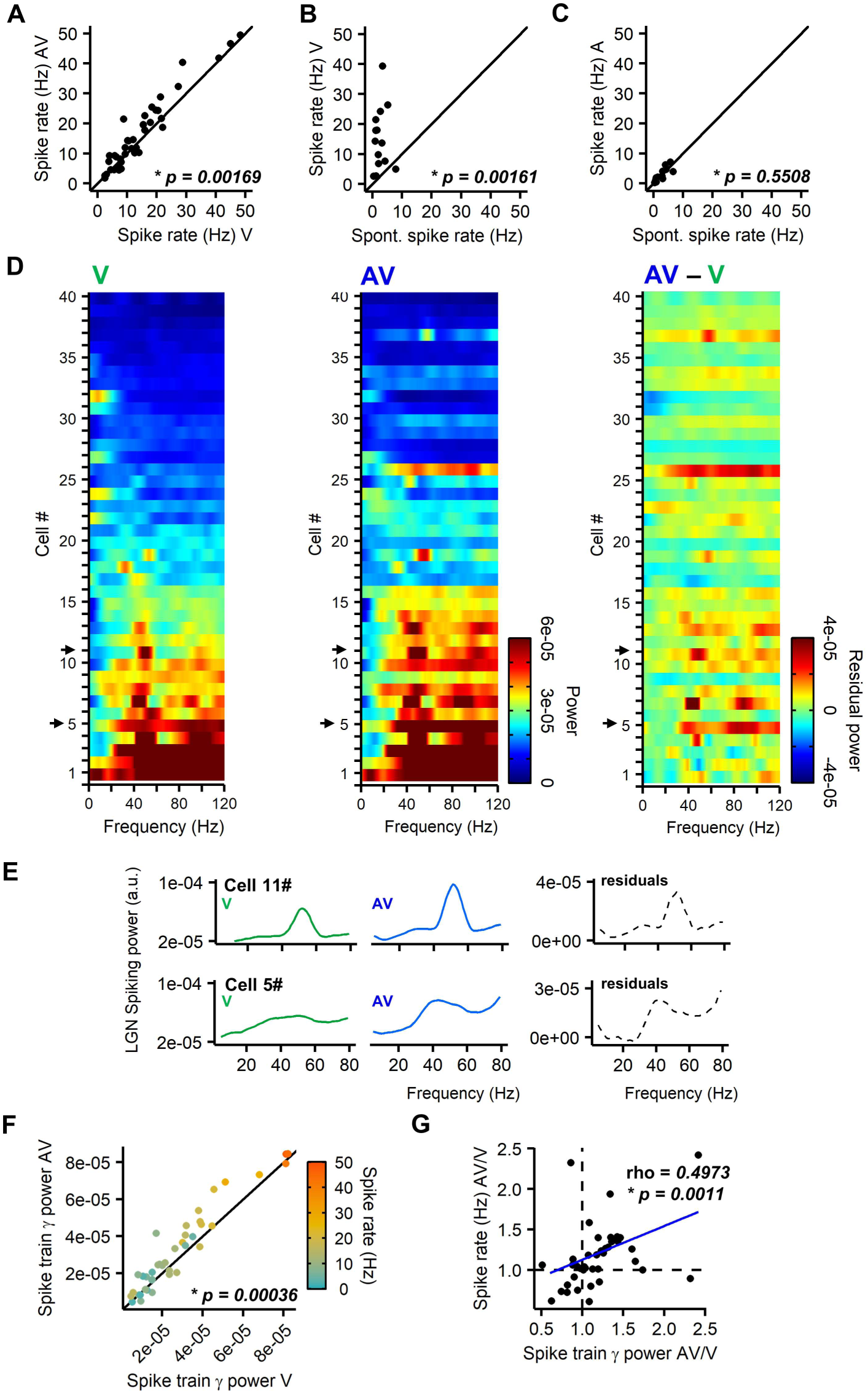
Audiovisual stimuli increase spike rate and spike train gamma power of LGN neurons. (**A**) Comparison of spiking rates of LGN neurons during visual (V) and audiovisual (AV) stimulation (n = 40, N = 10). **(B)** Comparison of LGN neurons spike rate during spontaneous activity and visual (V) stimulation (n = 14, N = 6). **(C)** Comparison of spiking rates of LGN neurons during spontaneous activity and auditory (A) stimulation (n = 14, N = 6). **(D)** Pseudocolor plot showing the spike train power spectrum (sorted by power spectral density) of the entire population of LGN neurons during visual (V, left), audiovisual condition (AV, middle) and their residuals (AV-V, right). **(E)** Spike train power spectra of two example neurons show an increase in the rhythmicity of their firing at gamma frequency in response to audiovisual stimuli. **(F)** Comparison of spike train gamma-band power of LGN neurons during visual (V) and audiovisual stimulation (AV); the color-code represents the spike rate of individual neurons. (A-C and F) Wilcoxon signed-rank test. **(G)** Relation between audiovisual-induced variation in spike train gamma-band power and spiking rate. The blue line corresponds to the linear regression and rho is the value of the Spearman’s rank correlation coefficient.

Although the majority of neurons recorded (55%, n = 22) increased their spiking rate (+3.9Hz (2.7)) in response to audiovisual stimuli, 20% (n = 8) showed a decrease (–2.6 Hz (1.4)) and the remaining 25% (n = 10) showed negligible or very little variation of their spiking rate (+0.07 Hz (0.34)). Figure S6A and S6B illustrate data from two example neurons. Furthermore, LGN-ON neurons (Figure 4B) did not exhibit cross-modal auditory response (Figure 4C), thus ruling out a potential contribution of sound alone in the increasing effect of audiovisual stimuli on LGN spiking.

To explore a potential connection between the audiovisual properties of the LGN and the increase of V1 NBG power in response to audiovisual stimuli (Figure 2 and 3), we examined the rhythmicity of LGN firing. Inspection of LGN spike train power spectra during visual-evoked activity revealed that ⁓50% of the recorded neurons exhibited rhythmical firing at frequencies falling within the gamma-band (Figure 4D). Spike train gamma-band power (40– to-80 Hz band power) was strongly correlated with the firing rate of individual neurons (Rho = 0.7424, *p* < 0.001; Figure S7). Audiovisual stimuli remarkably increased the spike train gamma-band power of most neurons exhibiting rhythmic gamma firing (*p* = 0.0003; Figure 4D-4F). Audiovisual-induced changes in spike rate and in spike train gamma power exhibited moderate positive covariance (Rho = 0.4973, *p*= 0.0011; Figure 4G). This observation indicates that when audiovisual stimuli cause a change in the LGN spike rate, there is likely to be a corresponding change in the gamma power of the spike train, and vice versa. Collectively, data from these experiments indicate that LGN-ON neurons exhibit audiovisual properties and suggest a thalamic origin of the V1 NBG audiovisual response.

## Discussion

Our finding that the power of V1 NBG is modulated in response to audiovisual stimuli suggests that corticocortical transfer of auditory information from A1 to V1 (Deneux et al., 2019; Garner et al., 2022; Ibrahim et al., 2016; Iurilli et al., 2012) may not be the sole source of audiovisual information integration in V1. This likelihood was confirmed by our observation that local pharmacological inhibition, or local optogenetically-mediated suppression of A1 auditory responses failed to interfere with the V1 NBG audiovisual response. This contrasts with prior studies that focused exclusively on direct corticocortical connections from A1 to V1 and served as the basis for current assumptions in the field that the sole source of auditory information transfer to V1 comes from A1 (Deneux et al., 2019; Garner et al., 2022; Ibrahim et al., 2016; Iurilli et al., 2012). The audiovisual properties of LGN that we identified, namely an increase of the visual-evoked response and gamma-synchronous firing, strongly support that the LGN constitutes a subcortical source for the transfer of audiovisual information to V1. Although our experiments did not yield direct empirical evidence as to the thalamic origin of the audiovisual effect that we detected at the cortical level, this interpretation is strongly supported by recent compelling evidence pointing to a direct causal link between V1 NBG and the firing of LGN ensembles (McAfee et al., 2018; Meneghetti et al., 2021; Saleem et al., 2017; Shin et al., 2023). Moreover, we are confident that our findings are clearly associated with visual NBG, given that the cortical oscillation we detected in V1 layer 4 exhibited a sharp peak of ⁓69 Hz and a narrow bandwidth (59-82 Hz) that never extended to the range of broadband gamma oscillations (30–90 Hz). The slightly higher frequency and wider bandwidth detected in our anesthetized mice compared to awake mice (Saleem et al., 2017) possibly derives from the permissive effect of the anesthetic on gamma oscillations (Imas and Hudetz, 2005).

We observed that sound pressure levels of 70-dB SPL, and higher, that occur coincidentally, during, or begin before (at least 100 ms), visual stimuli, all enhanced the power of V1 NBG. The integration of auditory inputs into the visual response of V1, despite temporal mismatches between these modalities, highlights the flexibility and adaptability of V1 in processing audiovisual information. Moreover, V1 NBG audiovisual response lingered after cessation of audiovisual stimuli, suggesting that bimodal audiovisual stimuli may transiently primed V1 towards a higher level of activation consistent with the creation of an electrophysiological substrate for the generation of an experience-dependent record (Galuske et al., 2019). This property raises the intriguing possibility that the audiovisual modulation of LGN-driven V1 NBG may play a particular role in the determination of subsequent cognitive associations (Fries, 2005, 2015; Salinas and Sejnowski, 2001). V1 NBG audiovisual responses were particularly potent following brief elevations in sound levels delivered in the middle of a visual oscillatory period. This audiovisual paradigm, used in the present study, highlighted that the V1 NBG audiovisual response unfolded as a bimodal sequence, beginning with an initial transient desynchronization followed by an enhanced level of synchrony. We postulate that this property arises through the combined activation/modulation of crossmodal corticocortical (A1-V1) (Deneux et al., 2019; Garner et al., 2022; Ibrahim et al., 2016; Iurilli et al., 2012) and audiovisual thalamocortical (LGN-V1) inputs, reported in the present study. The temporal organization of these two contrasting events is likely to derive from the varying durations in the activation/modulation of these two pathways. The former exhibits a short-lasting response to sound onset, whereas the latter reflects a sustained increase in thalamocortical synchrony (McAfee et al., 2018; Meneghetti et al., 2021; Saleem et al., 2017; Shine et al., 2023). At the functional level, this feature may first serve to promote momentary selective cortical processing of unexpected sound stimuli (Iurilli et al., 2012) and subsequently enhance visual processing of luminance information.

We propose that the LGN is an important source of audiovisual information for V1. The question arises, however, as to how this information is conveyed within the subcortical circuits and how the LGN acquires it. The absence of LGN crossmodal auditory response implies that its audiovisual properties arise from the specific temporally congruent bimodal interaction between visual and auditory inputs. Our initial intuition was that the tectogeniculate pathway, linking the superficial layers of the superior colliculus (sSC) to the LGN (Bickford et al., 2015), might contribute to this aspect by conveying audiovisual inputs, in addition to visual inputs (Ahmadlou et al., 2018). However, audiovisual responses in the SC are predominantly restricted to its deep layers (dSC) (Bednárová et al., 2018; Ito et al., 2018 and 2021; Meredith and Stein, 1986). The tectogeniculate pathway is, thus, unlikely to serve as a source of audiovisual information to the LGN. An alternative instigator of audiovisual information transmitted to the LGN may simply comprise stimulus-driven changes in arousal. In line with this possibility, a recent study has demonstrated that retinal outputs, which act as the primary upstream source of visual NBG (Koepsell et al., 2009; Neuenschwander and Singer, 1996; Storchi et al., 2017), are modulated by the level of arousal (Schröder et al. 2020). This form of modulation was evidenced in the SC (Schröder et al. 2020), but its occurrence in LGN and its sensitivity to audiovisual stimuli remains unknown. Arousal-mediated audiovisual processing by the LGN, in turn may be supported by reciprocal connections existing between the LGN and the thalamic reticular nucleus (TRN) (Halassa et al., 2014; McAlonan et al., 2008) that could support the temporal relationship we observed between sound onset and the increase in V1 NBG power. Moreover, the TRN was recently described as a powerful regulator of thalamocortical visual NBG (Hoseini et al., 2021). Nonetheless, our measures of pupil diameter in response to audiovisual stimuli did not reveal any discernible changes in arousal. Investigating these aspects more broadly will provide a more comprehensive understanding of how audiovisual information is transferred and integrated within the visual system.

In conclusion, our study demonstrate a new role for the LGN as an audiovisual integration and relay center and provides novel insights as to the sources of audiovisual information transfer to V1.

## Grants

This research was funded by grants from the Deutsche Forschungsgemeinschaft (DFG, German Research Foundation, www.dfg.de), SFB 874/A9 & B1 – project number: 122679504 to P.K. and D.M-V; SPP 2411 – project number: 520284247 to D.M-V. The funding organization had no role in study design, data collection and analysis, decision to publish, or preparation of the manuscript.

## Acknowledgments

The authors thanks Ann-Christin Ammann and Josephine Ansorge for technical support and Katja Reinhard (SISSA, Italy) for helpful discussion.

## Author contributions

Conceptualization, C.E.L., P.K., and D.M-V.; Methodology, C.E.L.; Experimentation, C.E.L.; Data analysis, C.E.L.; Writing-Original draft, C.E.L. and D.M-V.; Writing-Review & Editing, C.E.L. and D.M-V.

## Supplemental information

**Supplementary Figure 1.**
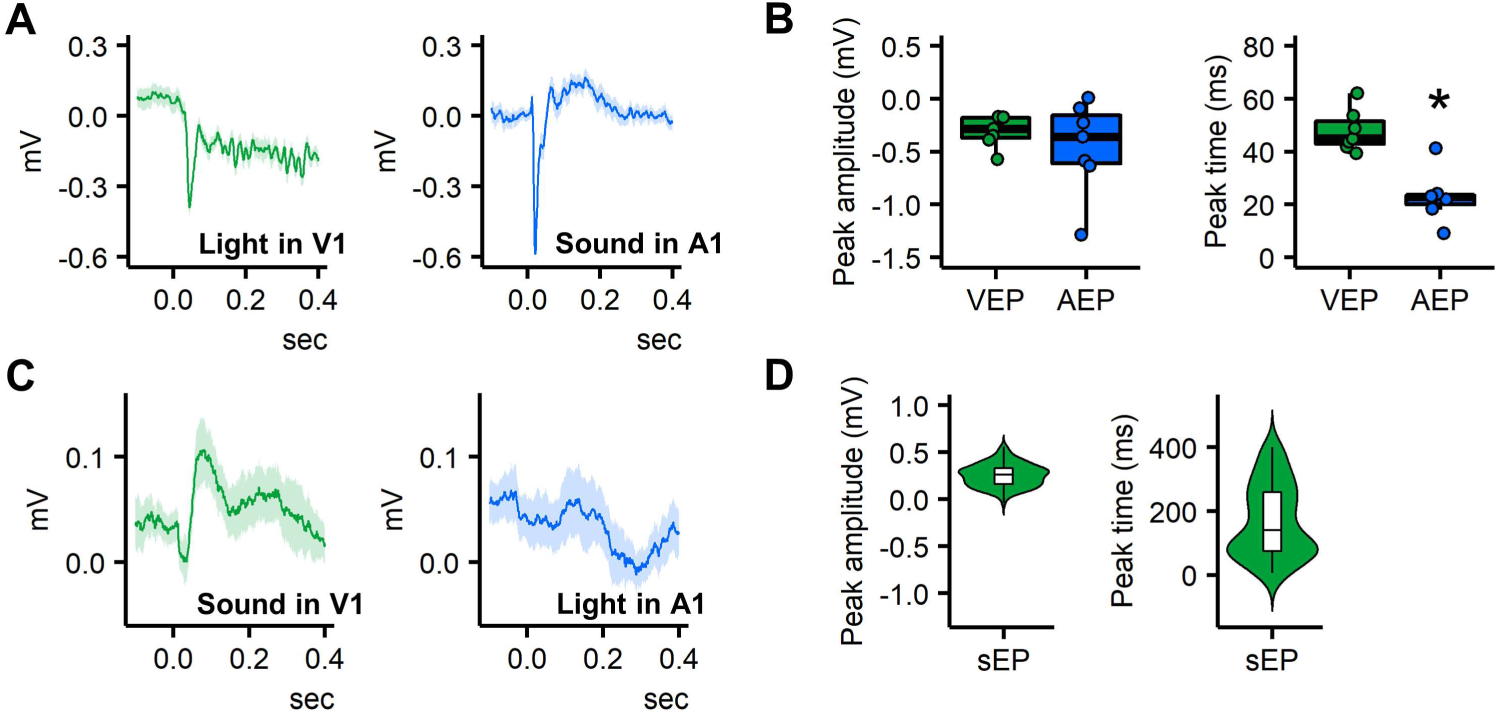
Response properties of V1 and A1 to unimodal and cross-modal sensory stimulation. **(A)** Examples of LFP waveform (average of 20 trials, data from the same animal) showing visual-evoked potential in V1 (VEP, left) and auditory-evoked potential in A1 (AEP, right). **(B)** Quantification of VEP and AEP peak amplitude and peak time (* *p* = 0.0156; Wilcoxon signed-rank test). **(C)** Grand mean V1 and A1 LFP waveform in response to cross-modal stimulation (average of 130 trials, N = 7 mice). Note that V1 LFP exhibits a cross-modal auditory response, which is manifested by a positive deflection. **(D)** Violin plots with boxplots overlay showing the distribution of peak amplitude and peak time of the sound-evoked potential in V1 (sEP).

**Supplementary Figure 2.**
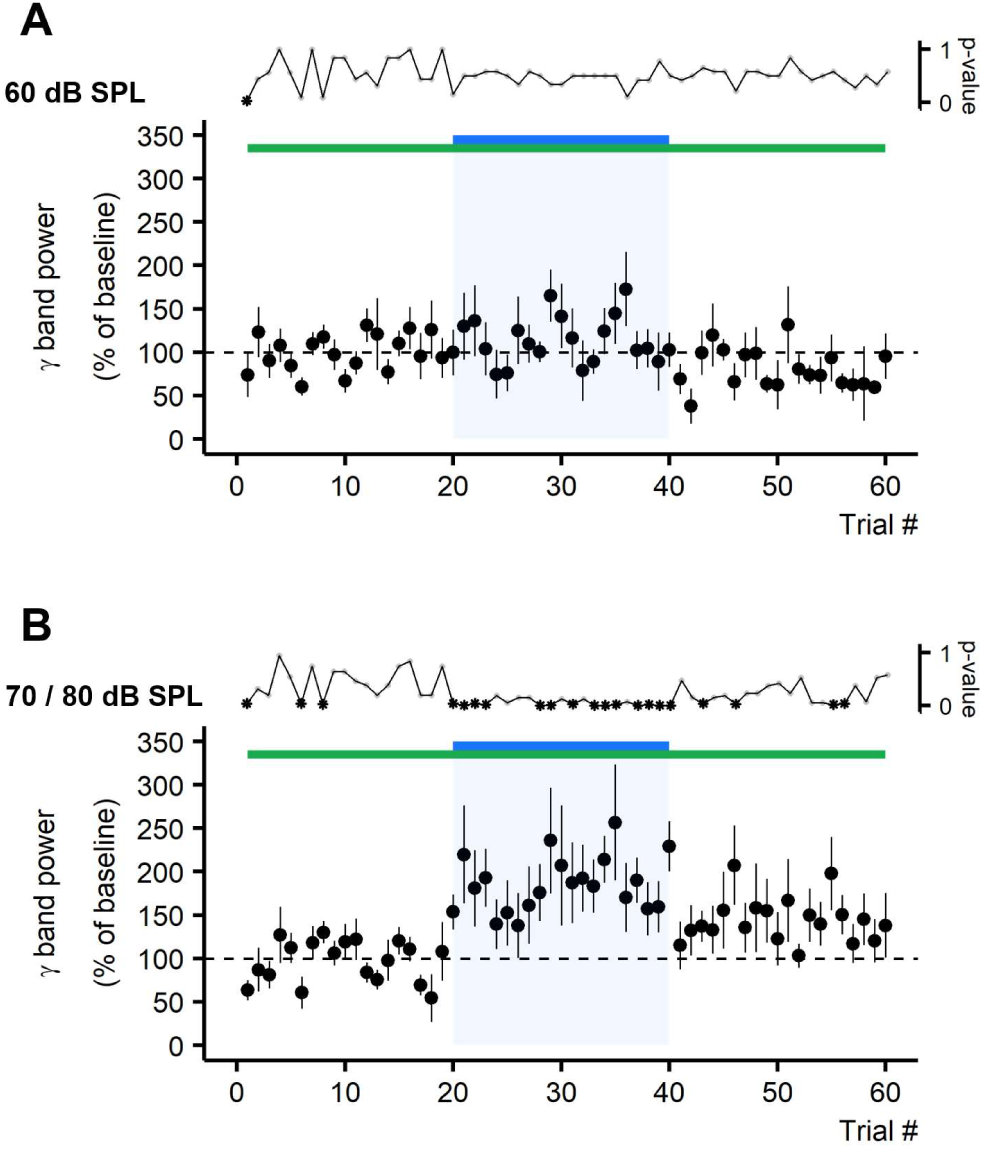
Temporal dynamic of V1 NBG power before, during, and after audiovisual stimuli. (**A-B**) Inter-trial variation in V1 NBG power (40-to-80 Hz band power) along three consecutive recording blocks comprising two blocks with visual stimuli alone interleaved by one block with audiovisual stimuli in which sound is delivered at 60–(A) or 70/80-dB SPL (B) (AV 60-, N = 6; AV 70/80-dB SPL, N = 8). Inter-trial time interval was 20 sec and trial duration was 6 sec. To increase the number of observations with post-AV monitoring, data from 70– and 80-dB SPL satisfying this criterion were pooled. (*\** indicate significant differences (*p*-value < 0.05) between each time point and the average of the first 20 sweeps, Wilcoxon signed-rank test)

**Supplementary Figure 3.**
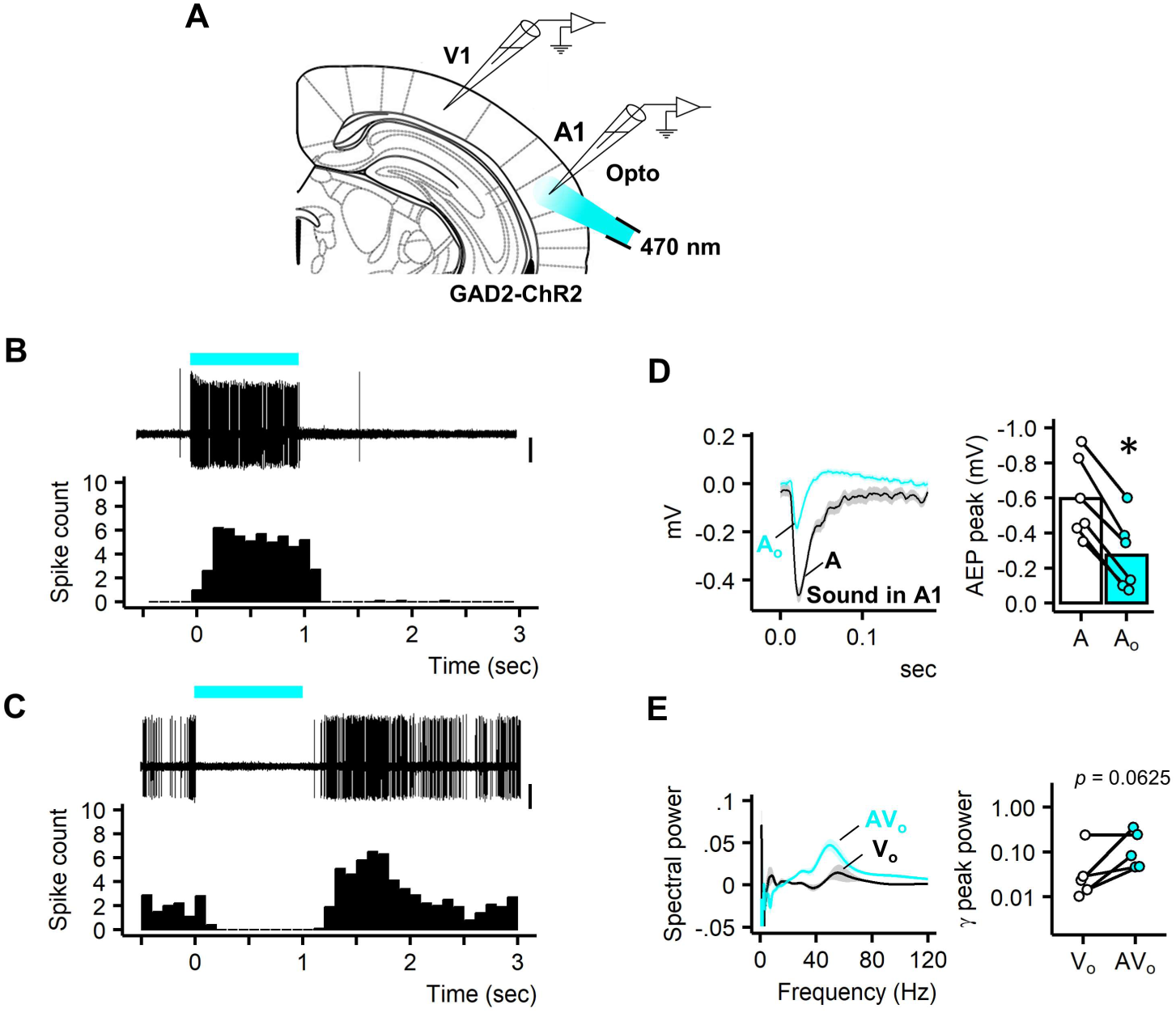
V1 NBG audiovisual response is unaffected in condition of A1 photo-inactivation. (**A**) Illustration of the setup used for experiments with photo-inactivation of A1 (Franklin and Paxinos, 2008; Plate 54). **(B)** Juxtacellular spike traces (overlay of 5 sweeps) showing the photo-activation of a putative Gad2-expressing neuron. Below is shown the peri-stimulus time histogram (average of 10 sweeps). **(C)** Juxtacellular spike traces (overlay of 5 sweeps) showing the inhibition of a putative excitatory neurons through photo-activation of Gad2-expresing neurons. Below is shown the peri-stimulus time histogram (average of 10 sweeps). Example data in (B) and (C) are from two different animals. Scale bars: 0.4 mV. **(D)** Average auditory response before (dark) and during (cyan) photo-activation of Gad2-positive neurons (average of 60 trials, N = 6; range of light intensity of photo-stimulation: 0.6 – 2.4 mW); quantification of the auditory response peak amplitude without and with photo-stimulation (N = 6; *p* = 0.0312, Wilcoxon signed-rank test). **(E)** Examples of power spectra showing V1 NBG during V and AV stimuli in condition of photo-inactivation of A1 (average of 20 trials, data from the same animal); quantification of NBG peak power (80-dB SPL, N = 5; *p* = 0.0625, Wilcoxon signed-rank test). Note that the power of V1 NBG increases in all experiments. In (D-E), the lower-case letter ‘o’ denotes the photo-stimulation condition.

**Supplementary Figure 4.**
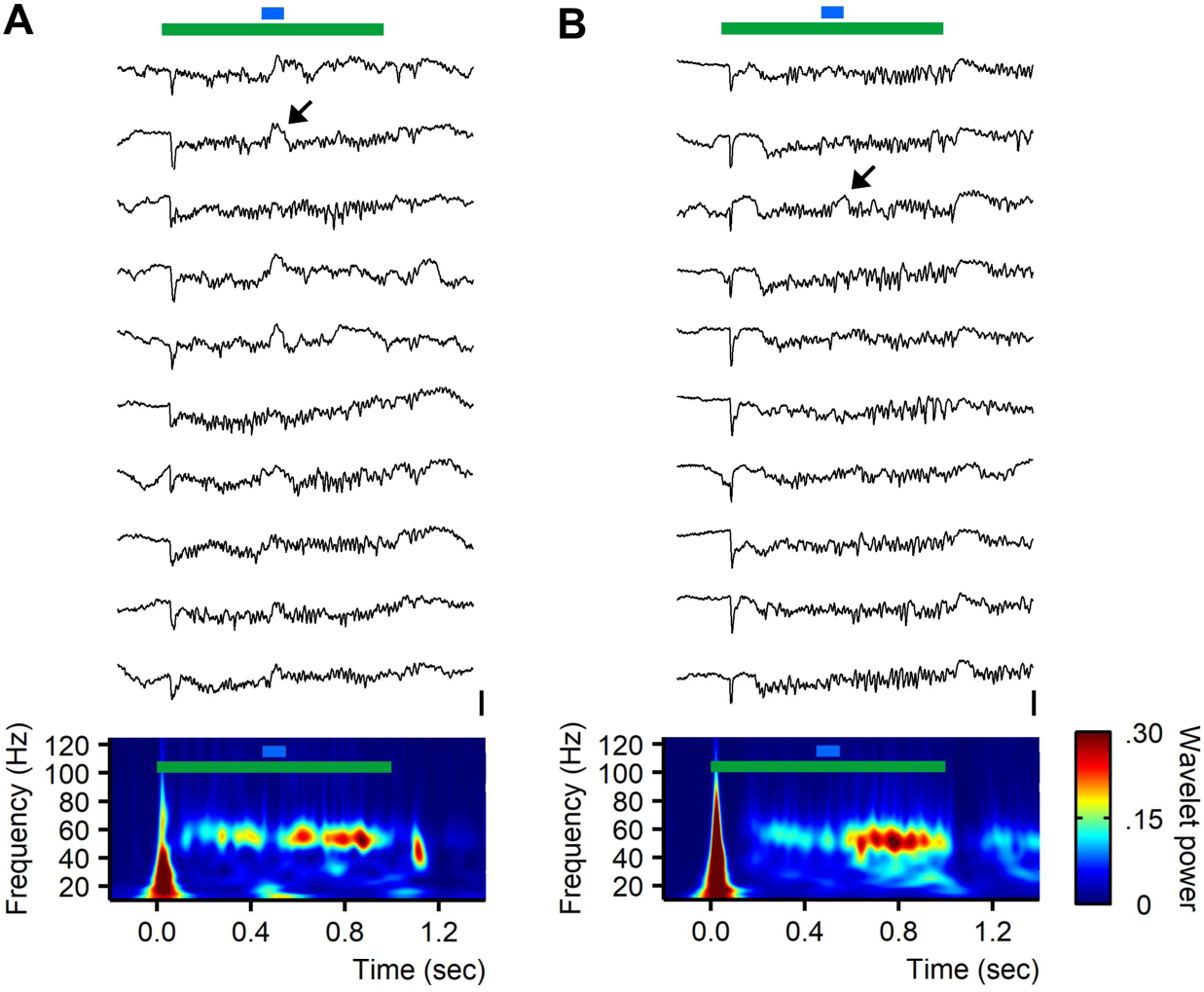
V1 NBG audiovisual response unfolds as a bimodal sequence. (**A-B**) Example data of ten consecutive trials for audiovisual stimuli with auditory stimulus presented in the middle of an ongoing V1 NBG (data from two independent experiments). Arrows are pointing to sound-induced up-ward deflection in V1 LFP. Scale bars: 0.6 mV. Bellow the traces is shown the corresponding wavelet power spectra (average of 20 trials).

**Supplementary Figure 5.**
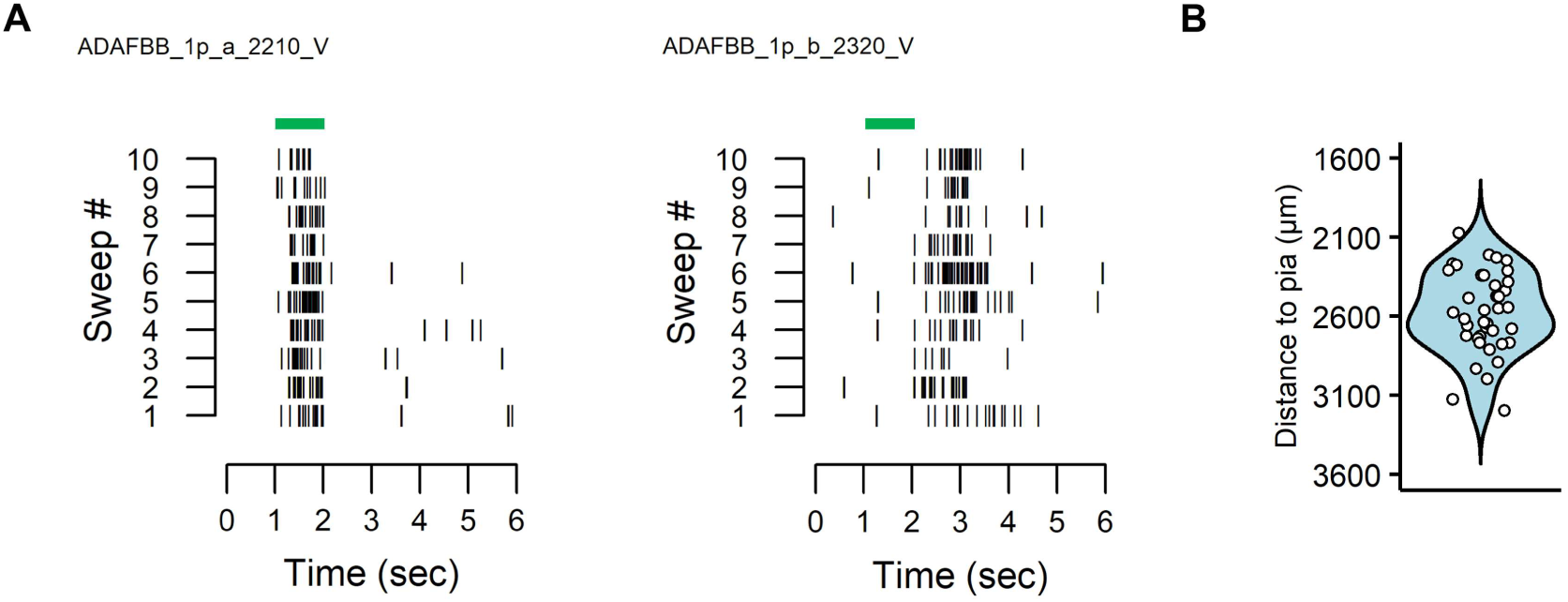
Recording of LGN neurons. (**A**) Example of LGN neurons (recorded in the same animal) activated either by switch-on (LGN-ON) or by switch-off (LGN-OFF) visual stimulation (data from ten consecutive trials). **(B)** Population depth distribution of LGN-ON neurons (n = 40, N = 10).

**Supplementary Figure 6.**
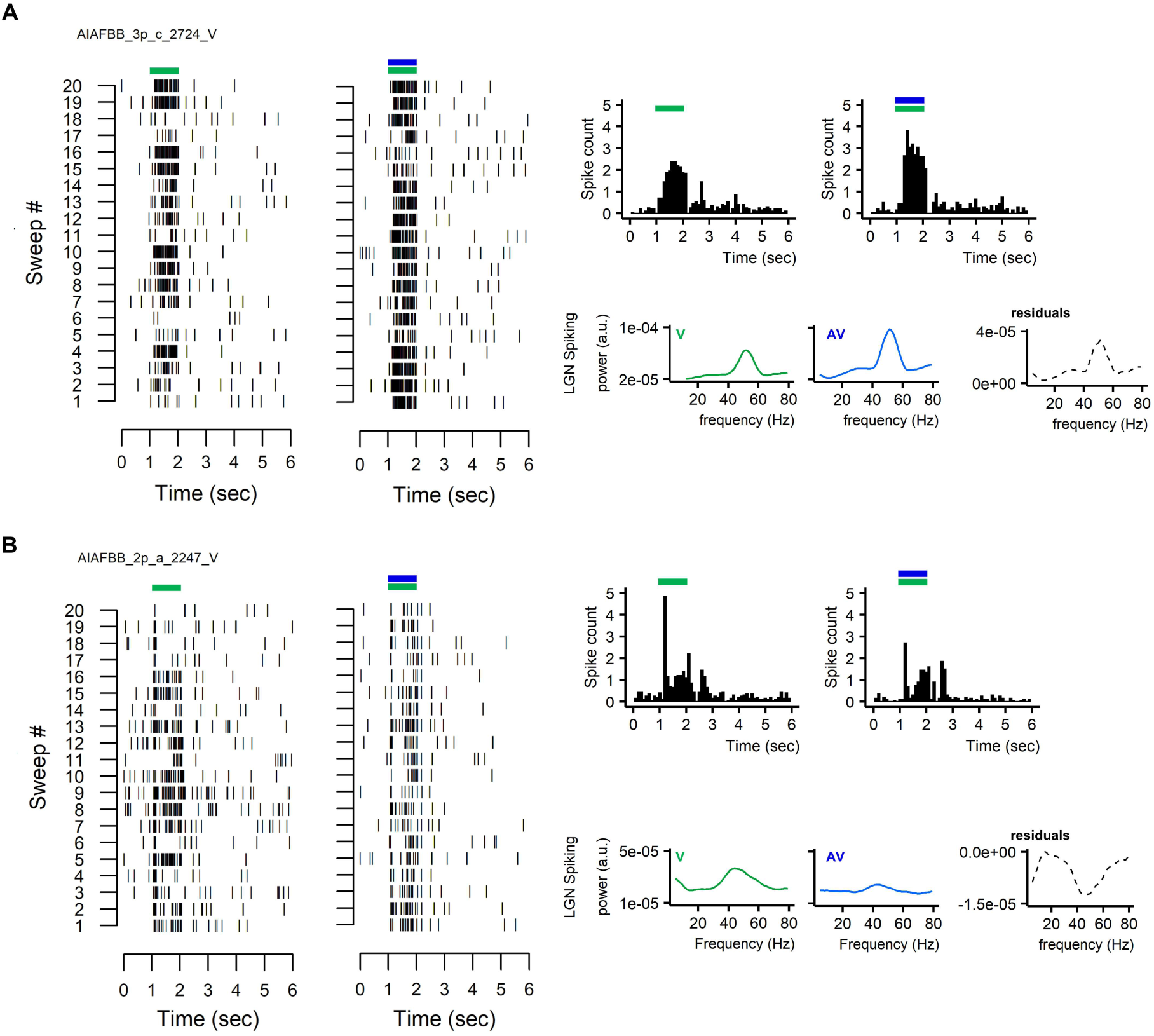
Audiovisual modulation of LGN activity: example neurons. (**A-B**) Spike raster plot, peristimulus time histogram (bin width: 10 ms) and spike train power spectra plus residuals for 20 consecutive trials either during visual or audiovisual stimulation for two example neurons; in (A) up-regulated and in (B) down-regulated by audiovisual stimuli. These two neurons were recorded in the same animal.

**Supplementary Figure 7.**
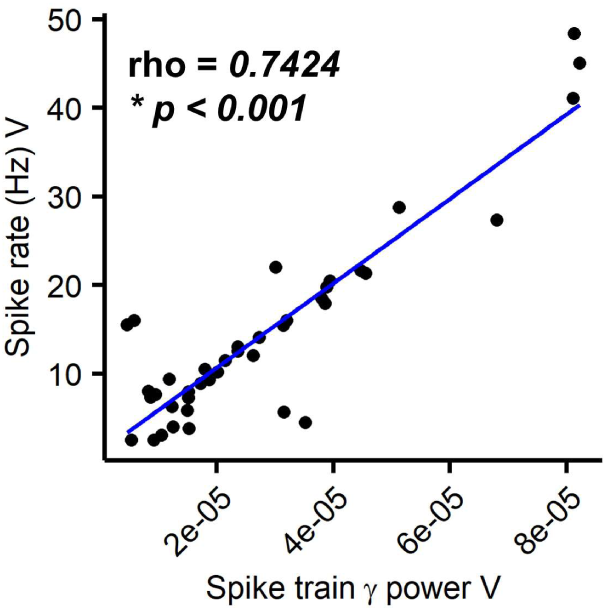
Relation between visual-evoked LGN spike rate and spike train γ power. The blue line corresponds to the linear regression and rho is the value of the Spearman’s rank correlation coefficient (n = 40, N = 10).

